# Exercise engages a mechanically activated astrocyte state linking muscle activity to hippocampal plasticity

**DOI:** 10.64898/2026.08.06.742872

**Authors:** Md Saddam Hossain Joy, Ilber Erden Manavbasi, Ki Yun Lee, Meghan Grace Connolly, Quinlan Valkyrie Glueck, Movviz Ahmed, Mohammad Adel Zayyad, Younes Haidari, Bashar Emon, James Song, Yash Deswal, Daniel Lee, Joseph P. Ritchie, Nora Sripraram, Justin S. Rhodes, M. Taher A. Saif

## Abstract

Physical exercise promotes brain health in part through muscle-derived factors that enter the brain, but how peripheral signals are translated into neural responses remain unclear. Here, we identify astrocyte contraction as a previously unrecognized physiological response to exercise signals that may contribute to adult hippocampal neurogenesis. In vivo, voluntary running rapidly induced nuclear localization of the mechanically sensitive transcriptional regulator Yes-associated protein (YAP), and increased non-muscle myosin II phosphorylation in hilar astrocytes in mice, consistent with acute contraction. Using an in vitro platform with ultrasensitive force sensors, we found that factors released by contracting skeletal muscles activated astrocytes which in turn increased contraction that was necessary and sufficient for their proliferation and expansion. Activated astrocytes subsequently released soluble factors that modulated neuronal network tension and promoted immature neuron abundance. These findings identify astrocyte contractility as a physiological transducer of exercise-derived muscle signals and establish cellular force generation as a potential mechanism regulating neuroplasticity.

## INTRODUCTION

It is well established that physical exercise is among the most important lifestyle factors that promote mental health and protect against cognitive decline. (*1–3*) In mice, voluntary exercise enhances adult hippocampal neurogenesis and synaptic plasticity, while improving learning and memory and attenuating age-related cognitive decline. (*4–6*) Although numerous circulating exercise-induced factors have been identified, (*7–13*) how peripheral muscle activity is translated into coordinated cellular responses within the hippocampus remains incompletely understood.

Neural function has traditionally been explained through electrical activity and biochemical signaling (*14*). However, cells also generate, transmit, and respond to mechanical forces that regulate proliferation, migration, differentiation, gene expression, and tissue organization throughout biology (*15–20*). In the nervous system, mechanical cues influence neural development, regeneration, and disease, and accumulating evidence indicates that neural cells actively sense their mechanical environment (*15*, *16*, *21–24*). Whether regulated cellular force generation also contributes to normal physiological signaling in the adult brain, however, remains largely unexplored.

Astrocytes are ideally suited to participate in mechanically regulated signaling. They possess integrins, a highly organized actomyosin cytoskeleton, and the mechanosensitive YAP/TAZ pathway, providing the molecular machinery required for force sensing and generation (*17–19*). In reactive astrocytes and glioma cells, activation of these pathways drives cytoskeletal remodeling, migration, and invasion, showing that astrocyte-lineage cells possess substantial force-generating capacity (*20*, *27–29*). Whether mature astrocytes engage regulated actomyosin contractility during normal physiology, and whether such mechanical responses contribute to hippocampal plasticity, remains unknown.

Exercise provides an ideal physiological context in which to address these questions. Astrocytes are increasingly recognized as essential mediators of exercise-induced hippocampal plasticity through their regulation of neurogenesis, synaptic function, and neuronal survival (*30–32*). Because astrocytes directly interface with both the cerebral vasculature and hippocampal neural circuits, they are ideally positioned to detect circulating exercise-induced signals and coordinate local plasticity. Moreover, within the dentate gyrus, astrocytes provide trophic, metabolic, and other paracrine signals that regulate the proliferation, survival, maturation, and integration of newly generated neurons (*32*, *33*). Consistent with this role, we previously demonstrated that muscle activity alone is sufficient to stimulate hippocampal astrocyte proliferation in vivo, and that conditioned media from contracting skeletal muscle enhances astrocyte proliferation, neuronal activity, and synaptogenesis in primary hippocampal cultures. These findings suggest that astrocytes function as intermediaries linking peripheral muscle activity to hippocampal plasticity, but the cellular mechanisms through which they transduce exercise-derived signals remain unknown.

Here, we tested the hypothesis that exercise engages a mechanically activated astrocyte state that contributes to hippocampal plasticity. We first examined mechanical signaling and activation of the astrocyte actomyosin machinery during voluntary running in vivo in mice. We then developed an in vitro platform integrating traction force microscopy, tunable extracellular matrix stiffness, and ultrasensitive force sensing to test causal relations between astrocyte contractility and exercise-induced plasticity. We show that voluntary exercise is associated with rapid activation of astrocyte mechanical signaling in vivo. In vitro, muscle-derived factors induced sustained, stiffness-dependent astrocyte contractility that was both necessary and sufficient to promote astrocyte expansion and activation. We further show that the contractility of the astrocytes affects their function in terms of regulating the contractility of mature hippocampal neurons and the abundance of immature neurons in primary culture. Together, these findings identify astrocyte mechanical contractility as a previously unrecognized component of the cellular response to exercise and support a model in which regulated cellular force generation contributes to communication between peripheral muscle activity and hippocampal plasticity.

## MATERIALS AND METHODS

### Animals

For *in vivo* analyses of YAP nuclear translocation and phosphorylated non-muscle myosin II regulatory light chain (pMLC) expression in hippocampal astrocytes, 8-week-old male C57BL/6J mice (n = 20) were housed either with or without access to running wheels, as indicated below. For preparation of primary muscle cell cultures used to generate muscle-conditioned media (CM), skeletal muscle was harvested from a total of six 4-week-old CD1 mice of both sexes. Primary hippocampal cultures for *in vitro* experiments were prepared from postnatal day 2 (P2) CD1 mice. All animal procedures were approved by the University of Illinois Institutional Animal Care and Use Committee (IACUC) and were conducted in accordance with National Institutes of Health guidelines.

### *In vivo* analysis of YAP nuclear translocation and pMLC expression in hippocampal astrocytes after wheel running

Mice were acclimated in groups of 3–4 for 2 weeks after arrival and then housed individually with (n = 10; runner) or without (n = 10; sedentary) a running wheel (9-inch diameter mounted in the cage) for 2 or 3 days (evenly split across groups). Wheel rotations were recorded continuously in 1-min increments. Throughout the study, mice were maintained on a 12:12 h light:dark cycle (lights off at 7:00 PM) with food and water available ad libitum.

Mice were euthanized by CO₂ asphyxiation between 9:00 and 10:00 PM during peak activity. Runner and sedentary mice were sampled in alternating order, and runners were euthanized regardless of whether they were actively running at the time of collection. Immediately after euthanasia while the heart was still beating, mice were transcardially perfused with chilled 0.9% saline to remove the blood. Brains were then rapidly removed, immersion-fixed overnight in 4% paraformaldehyde, cryoprotected in 30% sucrose, and sectioned coronally at 40 μm using a cryostat. Sections were stored in cryoprotectant at −20°C until immunohistochemistry. Three hippocampal sections per animal (rostral, middle, and caudal) were analyzed.

Free-floating sections were washed in TBS, blocked for 45 min in TBS containing 4% goat serum and 0.2% Triton X-100 (TBS-X+), and incubated overnight at 4°C with antibodies against GFAP (chicken, 1:1000) together with either YAP (rabbit, Cell Signaling cat#14074; 1:250) or phosphorylated non-muscle myosin II regulatory light chain (rabbit, Invitrogen cat# PA1-10004; 1:1000 dilution). After washing, sections were re-blocked for 30 min and incubated for 2 h at room temperature with goat anti-rabbit Alexa Fluor 647 (Jackson Immuno, 1:250) and goat anti-chicken Alexa Fluor 488 (Jackson Immuno, 1:250). Sections were counterstained with DAPI (1:1000), mounted onto gelatin-coated slides, and coverslipped using ProLong Gold Antifade reagent.

Confocal z-stacks were acquired using a Zeiss LSM 880 microscope equipped with a 40× oil objective (NA 1.2). Images were collected at 0.5 μm intervals using sequential acquisition of DAPI, GFAP, and either YAP or pMLC fluorescence. For YAP, images were collected bilaterally from both the dentate gyrus hilus and molecular layer. For pMLC, images were collected bilaterally from the dentate gyrus hilus only. Pinhole size was optimized independently for each fluorescence channel to maintain accurate spatial registration throughout the z-stack.

Image analysis was performed in IMARIS x64 (version 10.0.1). DAPI staining was used to generate nuclear surfaces (Surface 1), and the GFAP channel was masked within these nuclear volumes to isolate cytoplasmic GFAP signal. Astrocyte surfaces (Surface 2) were then generated from the masked GFAP channel using an intensity threshold that maximized astrocyte coverage while minimizing object fusion. Astrocytes were identified as GFAP-positive surfaces associated with intact DAPI-defined nuclei.

For YAP analysis, nuclear and cytoplasmic YAP fluorescence intensities were quantified separately for each astrocyte, normalized to their respective compartment volumes, and expressed as a nuclear-to-cytoplasmic YAP ratio. Thirty astrocytes per hippocampal region (10 cells per section from each of three sections) were analyzed for each animal, and the mean ratio was used as the representative value. The sample size of 30 cells per region was determined empirically based on stabilization of the running mean.

For pMLC analysis, phosphorylated myosin fluorescence intensity was quantified for each astrocyte, normalized to astrocyte volume, and averaged across all eligible hilar astrocytes within each animal to generate a single representative value for statistical analysis.

### Isolation and culture of primary skeletal muscle cells

Primary mouse skeletal muscle cells were harvested following our previously published protocol(*34*). After euthanasia, the hindlimbs were carefully removed, and the surrounding skin and connective tissue were peeled back to expose the underlying muscles. The isolated muscle tissues were immediately placed in ice-cold PBS (Corning) to prevent degradation and maintain tissue viability. Using dissection scissors, the tissues were finely minced into small fragments to facilitate subsequent enzymatic digestion.

The minced muscle tissues were digested in a solution composed of DMEM supplemented with 2.5% HEPES, 1% GlutaMAX (Gibco), 1% Penicillin-Streptomycin (Lonza), 400 U/ml collagenase (Worthington), and 2.4 U/ml dispase (Sigma). The tissues were incubated in this digestion media at 37°C for 30 minutes with occasional gentle agitation to enhance enzymatic activity. Following the initial digestion, the partially dissociated tissues were triturated using a sterile pipette in 0.25% trypsin (Gibco) to further break down remaining cell clusters. The resulting suspension was passed sequentially through 70-µm and 40-µm cell strainers to remove debris and obtain a homogeneous cell population.

To improve the purity of myoblasts and reduce fibroblast contamination, a pre-plating technique was applied. Dissociated cells were transferred to uncoated tissue culture flasks and incubated at 37°C for 3 hours to allow fibroblasts and other adherent cells to attach to the flask surface. The non-adherent, floating cells, enriched in myoblasts, were collected and seeded onto culture dishes coated with 0.1 mg/ml Matrigel (Corning) to support cell adhesion and growth at a density of 0.04 million cells/cm² (*35*). The cells were maintained in muscle growth media, which consisted of Ham’s F-10 Nutrient Mix supplemented with 20% fetal bovine serum (FBS), 1% GlutaMAX, 1% MEM Non-Essential Amino Acids (NEAA) (all from Gibco), 1% Penicillin-Streptomycin, and 0.5% chick embryo extract (US Biological). To promote cell proliferation, the media was further enriched with ice-cold 10 ng/ml basic fibroblast growth factor (bFGF), 20 µM forskolin, and 100 µM IBMX (all from Sigma).

The cells were kept below 70–80% confluency to ensure optimal growth and avoid contact inhibition. Once the culture reached 70–80% confluency, the growth media was replaced with muscle differentiation media to initiate myotube formation. The differentiation media comprised a 1:1 mixture of DMEM and Ham’s F-12 Nutrient Mix, supplemented with 10% horse serum, 1% GlutaMAX, and 1% Penicillin-Streptomycin (all from Gibco). Myotube maturation was monitored, and when contractions were observed, the media was replaced with pre-muscle-conditioned media (pre-MCM). The pre-MCM consisted of Advanced DMEM/F-12 supplemented with 1% GlutaMAX and 1% Penicillin-Streptomycin, providing an optimal environment for continued myotube functionality. Pre-MCM is also used in cell culture described in the section Hippocampal cell culture and collection of CM, RM, and AST-CM described later.

### Hippocampal dissection and cell isolation

Hippocampal dissection was performed as described in(*36*). For each experiment, the hippocampi of 5 to 15 pups were used. After decapitation, the brains were quickly removed from the skulls and transferred to cold Hibernate-E media (Gibco) on ice to preserve cellular integrity and viability during dissection. Under a stereomicroscope, the hippocampus was carefully isolated from other brain regions using fine sharp tweezers. Care was taken to avoid contamination with non-hippocampal tissues. Once isolated, the hippocampi were transferred into fresh cold Hibernate-E media, where they were finely minced into small fragments using sharp dissection scissors to facilitate enzymatic digestion.

The minced tissues were subjected to two consecutive digestion steps in 2 mg/ml papain (Sigma) solution. Each digestion step lasted 30 minutes and was carried out at 37°C with gentle agitation to ensure uniform exposure of the tissues to the enzyme. Between digestion steps, the tissues were gently resuspended to enhance dissociation. Following enzymatic treatment, the tissues were further dissociated mechanically using a fire-polished glass pipette. This step was performed with care to break apart remaining tissue fragments while minimizing mechanical stress on the cells.

The resulting cell suspension was then filtered sequentially through 70-µm and 40-µm cell strainers to remove undigested tissue debris and obtain a homogeneous population of single cells. The filtered cells were collected in fresh Hibernate-E media and processed further for downstream experiments.

### Hippocampal cell culture and collection of CM, RM, and AST-CM

Hippocampal neurons and astrocytes were cultured using specialized media formulations tailored to the specific experiment. These included RM (regular media as control), CM (muscle-conditioned media), astrocyte media conditioned by RM (Ast-RM), and astrocyte media conditioned by CM (Ast-CM). The collection of CM, RM, Ast-CM, and Ast-RM was performed following our previously published protocol (*34*) with slight modifications described below.

Generation of CM and RM: Following our previous protocol (*34*), mature spontaneously contracting primary myotubes were switched to Advanced DMEM/F-12 supplemented with 1% GlutaMAX and 1% Penicillin-Streptomycin. Media conditioned by contracting myotubes was collected every 24 h for 8 days, filtered (0.22 μm), aliquoted, and stored at −80°C. Control media (RM) consisted of the identical medium incubated for the same duration under identical conditions but without muscle cells. Before use, CM and RM were each diluted 1:1 with Neurobasal medium and supplemented with 10% horse serum, 1% GlutaMAX, and 1% penicillin-streptomycin.

Generation of Ast-CM and Ast-RM: Primary hippocampal astrocytes were cultured in basal medium (1:1 Advanced DMEM/F-12:Neurobasal supplemented with 10% horse serum, 1% GlutaMAX, and 1% penicillin-streptomycin). At approximately 70% confluence, astrocytes were exposed to either CM or RM for 24 h. The media were then removed, cells were washed, and fresh basal medium was added for an additional 24 h. This second medium was collected, filtered (0.22 μm), and stored at −80°C as Ast-CM or Ast-RM, respectively. This two-step protocol ensured that the collected media contained astrocyte-secreted factors released in response to prior CM or RM stimulation rather than residual muscle-conditioned media.

### Substrate functionalization for astrocyte culture

Astrocytes were cultured on polyacrylamide (PA) gel substrates with elastic modulus 1, 5 and 10 kPa(*37*), or on glass substrate. PA gel substrates were functionalized with fibronectin for astrocyte adhesion, following the protocol detailed in (*37*). Glass substrate was treated with 0.1mg/mL Poly-L-Lysine (PLL) overnight for surface functionalization and astrocyte adhesion. Soft PA gel substrates were used to modulate astrocyte force and to measure the forces using Traction Force Microscopy (TFM)(*37*).

### Glial Treatment

To measure the force of neuronal tissue without the influence of glial cells, a glial cell suppression protocol was implemented following the method described in(*38*). On day 1, the sample was treated with a cocktail of 20 μM fluorodeoxyuridine (FUDR) (MP Biomedicals), 20 μM uridine, and 0.5 μM Ara-C (all from Sigma) for 72 h to suppress the glial cells from the tissue. After 72 h, two-thirds of the cocktail solution was replaced with neuron culture maintenance media. The cultures were then maintained by re-feeding every 3 d with neuron culture maintenance media, with half of the media being replaced with fresh maintenance media.

### Experimental setup of force sensors and 3D neuronal tissue formation

The experimental setup of the force sensors was carried out following our published protocol(*39*, *40*) with slight modifications to optimize neural tissue formation and measurement precision. A thin gelatin layer was first formed in the well of a glass-bottom Petri dish to serve as a sacrificial layer. The force sensors were then positioned on this gelatin layer, with their base attached to a double-sided adhesive tape to ensure stability. The space between the sensor beams was carefully filled with gelatin, which provided structural support during tissue formation and helped eliminate air bubbles that could interfere with the sensor’s force measurements. The gelatin was melted at 37°C and thoroughly washed out after the tissue was formed to leave a clean and functional sensor.

The extracellular matrix (ECM) mixture was freshly prepared on ice to ensure optimal conditions for tissue formation. Type-I collagen from rat tail (Corning) was neutralized by sequentially adding 1N sodium hydroxide, 10X PBS, and deionized (DI) water to adjust the pH. Neutralized collagen was then combined with Matrigel (Corning) to achieve a final concentration of 1 mg/ml for each component. This ECM solution, acting as the tissue precursor, was stored on ice and kept in a 4°C chamber until used for the experiment to maintain its integrity and prevent premature polymerization.

Detailed illustration of the 3D neuronal tissue formation process between the grips of the force sensor is described in our previous papers(*23*, *39*, *40*). To maintain a cell-free region in the middle of the sensor, a high-density cell suspension (∼10 million cells/ml) of rat hippocampal neurons was first pipetted directly onto the grips of the force sensor. Capillary tension was typically sufficient to draw the cell suspension into the grips while displacing any trapped air. For cases where air bubbles persisted, low-pressure vacuuming was applied for 30–40 seconds using a vacuum desiccator to ensure complete removal of air. Any residual cell suspension in the middle region between the grips was carefully removed using a wiping technique to avoid contamination of the cell-free area.

After ensuring the cell suspension was confined to the sensor grips, the ECM solution (devoid of cells) was pipetted onto the grips and into the space between them, forming a capillary bridge. This setup was left at room temperature for 40–50 minutes to allow the ECM solution to polymerize and create a stable 3D tissue structure. The resulting tissue consisted of neuronal populations anchored at the two ends of the sensor grips, with a central ECM-only region between them. This design was intended to promote neurite extension into the ECM-filled space, facilitating synapse formation.

The contractile forces generated by the neurites were transmitted to the sensor beams, enabling precise measurement of neuronal activity and tissue dynamics. The setup ensured the integrity of the tissue and minimized artifacts, allowing for accurate and reproducible force measurements.

### Immunofluorescence and image analysis of primary hippocampal cell cultures

For immunofluorescence imaging of the primary hippocampal cell cultures, samples were first fixed in 4% paraformaldehyde (PFA) prepared in phosphate-buffered saline (PBS) for 1 hour at room temperature to preserve cellular structures. After fixation, samples were permeabilized using 0.2% Triton X-100 in PBS for 10 minutes to allow antibody access to intracellular targets. To minimize non-specific antibody binding, samples were incubated in a blocking solution consisting of 2.5% bovine serum albumin (BSA) and 2% normal goat serum (NGS) in PBS for 1 hour at room temperature.

Following blocking, samples were incubated overnight at 4°C with corresponding primary antibodies diluted in blocking solution. The following antibodies were used for the in-vitro experiments: Rabbit anti-YAP, Invitrogen cat# 13584-1-AP, 1:100 dilution; Chicken anti-GFAP, Invitrogen cat# PA1-10004, 1:1000 dilution; Chicken anti-MAP2, Abcam cat# ab5392, 1:10000 dilution; Rabbit anti-DCX, Abcam cat# ab18723, 1:500 dilution. After primary antibody incubation, samples were washed thoroughly with PBS five times, with each wash lasting 5 minutes. Next, corresponding secondary antibodies conjugated to fluorescent dyes were added, and the samples were incubated for 2 hours at room temperature in the dark. The secondary antibodies were: Goat anti-Chicken AF488, Abcam cat# ab150173, 1:1000 dilution; Goat anti-Rabbit AF488, Abcam cat# ab150081, 1:1000 dilution; Goat anti-Chicken AF568, Abcam cat# ab175711, 1:1000 dilution. To stain cell nuclei, samples were incubated with DAPI (Invitrogen cat# D1306; 1:1000 dilution) for 10 minutes, followed by three PBS washes to remove excess dye.

Image acquisition was performed using a confocal laser scanning microscope (LSM 710, Carl Zeiss AG, Oberkochen, Germany) equipped with an EC Plan-Neofluar 20×/ 40x/ 63x objective lens. Confocal z-stack images were acquired at appropriate excitation/emission wavelengths for each fluorophore. Image processing was carried out using ImageJ (NIH, Bethesda, MD, USA), and IMARIS software (version 9.6.0, Bitplane AG, Zurich, Switzerland).

### Brightfield time-lapse imaging and sensor displacement analysis

Brightfield time-lapse imaging of the tissues and force sensors was carried out using an inverted optical microscope (Olympus IX81) equipped with a 20× objective lens (Olympus America Inc., Center Valley, PA). The microscope was mounted on a vibration isolation table (Newport Corporation, Irvine, CA) to minimize external disturbances during imaging. An environmental chamber enclosing the microscope stage maintained physiological conditions: 37°C temperature, 5% CO₂, and 70% humidity, ensuring the health and activity of the cultured tissues during live imaging.

Images were captured using a Neo sCMOS camera (Andor Technology, Belfast, Northern Ireland) with a resolution of 1392 × 1040 pixels and a pixel size of 167 nm. Time-lapse images were used to monitor tissue dynamics and the displacement of force sensor gauges.

Sensor displacement was quantified using the template matching plugin in ImageJ, which allows for sub-pixel resolution tracking of the gauge tips. This method enabled accurate measurement of tissue-generated contractile forces based on the deflection of the sensor beams.

### Statistical Analysis

All statistical analyses were performed using R (version 4.4.0). *In vivo* YAP ratio was compared between runner and sedentary groups using a two-tailed unpaired t-test. For evaluating correlations between YAP ratio and pMLC concentrations with running levels, simple linear regressions were used. *In vitro* data were analyzed using two-way analysis of variance (ANOVA), with stiffness of the media (e.g., 1, 5, 10 kpa) and media type (regular media, RM, or conditioned media, CM), followed by Least Significant Difference post hoc tests to determine pairwise differences. If residuals displayed skewness outside the range of -1 and 1 and kurtosis outside -2 and 2, then the data were analyzed using a non-parametric permutation test using the permuco package (aovperm function). A p-value of less than 0.05 (P < 0.05) was considered statistically significant for all analyses.

## RESULTS

### Exercise rapidly activates astrocyte mechanical signaling in vivo in mice

We used two separate complementary measures to determine whether voluntary exercise engages astrocyte mechanical signaling in vivo. First, we quantified nuclear versus cytoplasmic localization of the mechanically sensitive transcriptional regulator YAP (*18*). YAP is known to translocate from the cytoplasm to the nucleus in response to increased cellular contractility, thereby increasing the ratio of nuclear to cytoplasmic YAP concentrations. This mechanically induced YAP translocation has been attributed to deformation of the nucleus due to intercellular forces and can occur on a timescale of minutes (*41*, *42*). Second, we measured the concentration of phosphorylated non-muscle myosin II regulatory light chain (pMLC), a marker of actomyosin activation (*43*, *44*), that drives cellular contractility. Analyses focused on astrocytes within the dentate gyrus hilus, the principal neurogenic niche, and the adjacent molecular layer.

Three-dimensional Imaris reconstructions of GFAP-positive astrocytes were used to quantify YAP ratios in individual cells (Figs. 1A–D). Mean hilar YAP ratios did not differ significantly between sedentary and runner groups (Fig. 1E). However, runners showed substantially greater inter-individual variability (Bartlett’s test, P = 0.0044), suggesting that astrocyte YAP localization changes dynamically during exercise rather than remaining persistently elevated.

**Figure 1.**
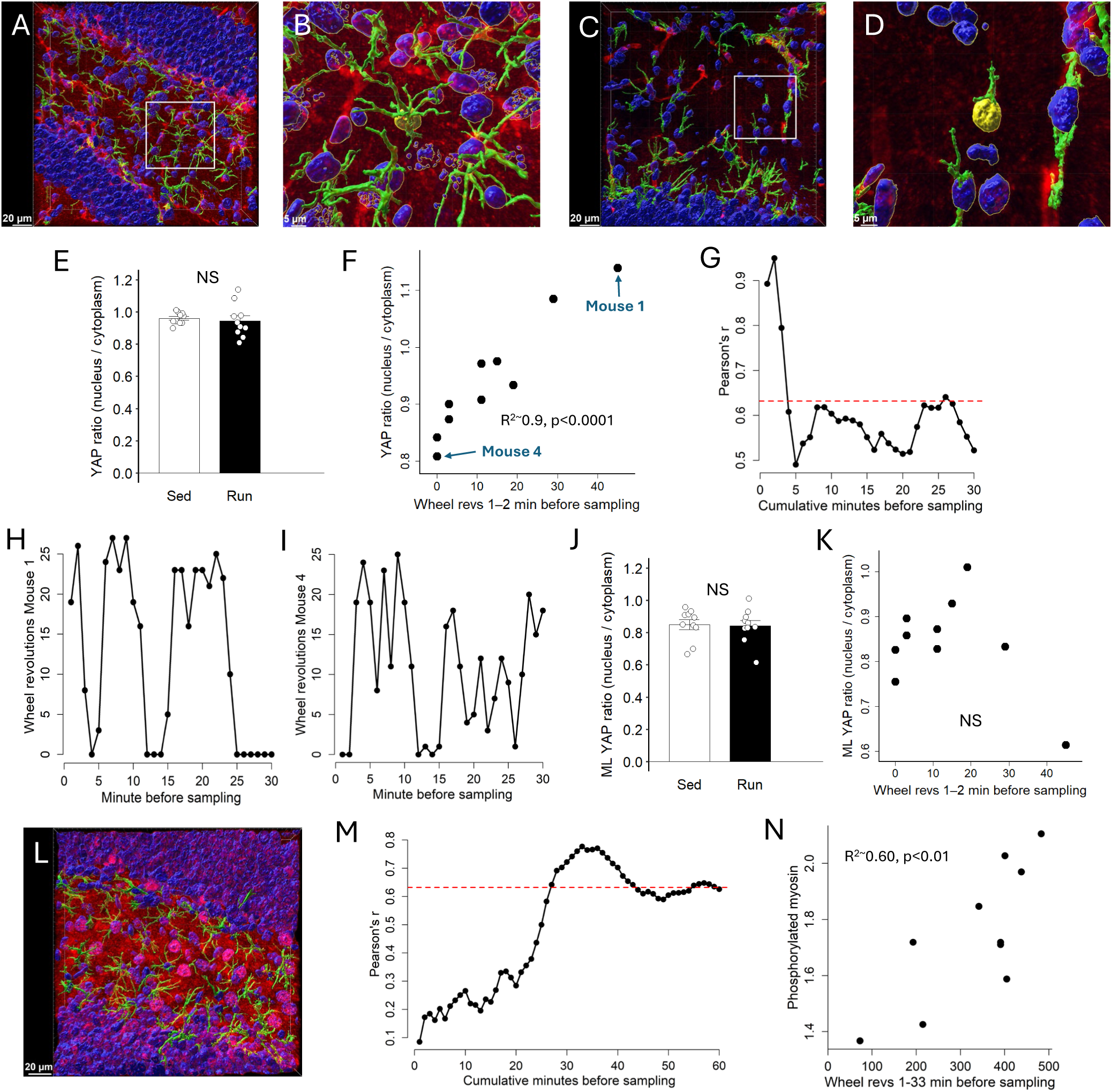
Running behavior dynamically engages mechanotransductive signaling in hilar astrocytes *in vivo*. (A) IMARIS reconstructed Z-stack image through the dentate gyrus hilus region showing DAPI (nuclei) in blue, GFAP (marks astrocytes) in green, and YAP in red. Scale bar: 20 µm. (B) Zoomed in image showing a hilar astrocyte eligible for data acquisition. The nucleus is highlighted in yellow and is surrounded by the cytoplasm highlighted in green. Scale bar: 5 µm. (C) Same as A through the molecular layer of the dentate gyrus. Scale bar: 20 µm. (D) Same as B for the molecular layer. Scale bar: 5 µm. (E) Mean +/- SE YAP ratio in hilar astrocytes shown for sedentary (Sed) and Runners (Run). No differences were detected between groups, runners displayed greater variation. (F) YAP ratio in hilar astrocytes plotted against wheel revolutions within the first 2 minutes before the runner was sampled. A strong positive correlation was observed. (G) Pearson’s correlations (r) between running activity and the astrocytic hilar nuclear-to-cytoplasmic YAP ratio are plotted as a function of the cumulative time window used to quantify running. The strongest relationships occur when running is measured over the first few minutes immediately preceding tissue collection, with correlations diminishing as longer durations are included. Correlations above the red dashed line are statistically significant p<0.05. (H) Wheel revolutions per minute for Mouse 1 (shown in panel F) plotted in 1-min bins across the 30 min preceding tissue collection. Each point represents the number of wheel revolutions during a single minute before sampling, illustrating the intermittent pattern of voluntary running, and active running immediately prior to collection. (I) Same as H except for Mouse 4 (shown in panel F), that was inactive immediately prior to collection. (J-K) same as E, F except for molecular layer astrocytes. No differences were detected between groups and no correlation with wheel revolutions was observed. (L) IMARIS reconstructed Z-stack image through the dentate gyrus hilus region showing DAPI (nuclei) in blue, GFAP (marks astrocytes) in green, and pMLC in red. Scale bar: 20 µm. (M) Same as G for astrocytic hilar pMLC intensity. The strongest relationships occur when running is measured between 30-38 min preceding tissue collection. (N) pMLC intensity in hilar astrocytes plotted against wheel revolutions within the first 33 minutes before the runner was sampled. A strong positive correlation was observed.

To determine the source of this variability in running mice and to assess whether YAP localization reflected recent locomotor activity, we correlated nuclear YAP localization with running behavior preceding tissue collection. Hilar astrocytic YAP localization was strongly associated with running during the immediately preceding 1–2 min (R² ≈ 0.9, *P* < 0.001; Fig. 1F). Animals collected during an active running bout exhibited robust YAP nuclear accumulation, whereas animals sampled shortly after running showed markedly lower nuclear localization. This relationship progressively weakened as the behavioral integration window increased (Figs. 1G), indicating that astrocytic YAP activation closely tracks recent locomotor activity rather than cumulative exercise. Consistent with the intermittent nature of voluntary running, animals with high nuclear YAP localization did not necessarily accumulate greater total running distance than animals with low YAP ratios (Figs. 1H,I). No association between running behavior and astrocytic YAP localization was observed within the molecular layer (Figs. 1J,K), indicating regional specificity of this response.

Because YAP reports mechanical signaling rather than force generation directly, we next examined activation of the astrocyte actomyosin machinery by quantifying pMLC immunofluorescence within hilar GFAP-positive astrocytes (Fig. 1L). Cumulative wheel running was calculated over progressively increasing intervals (1–60 min) before tissue collection and compared with astrocytic pMLC intensity. Significant positive correlations emerged after approximately 27 min of cumulative running and were strongest for intervals spanning 30–38 min before tissue collection (Fig. 1M). An example using cumulative running during the preceding 33 min is shown in Fig. 1N (R² ≈ 0.6, *P* = 0.008). In contrast to the minute-scale dynamics of YAP localization, pMLC was most closely associated with sustained running behavior over tens of minutes.

Together, these findings indicate that voluntary exercise is accompanied by rapid activation of astrocyte mechanical signaling followed by progressive engagement of the actomyosin machinery within the hilar neurogenic niche.

### Muscle-derived signals induce sustained astrocyte contractility in a stiffness-dependent manner

Motivated by the observations above that voluntary exercise activates astrocyte mechanical signaling in vivo, we next asked whether muscle-derived biochemical signals are sufficient to induce astrocyte contractility in vitro. We used our previously established muscle-to-hippocampus signaling model, in which conditioned media (CM) from spontaneously contracting primary skeletal myotubes is applied to primary hippocampal astrocytes (*34*). Astrocyte contractility was quantified by traction force microscopy (TFM) on fibronectin-coated polyacrylamide hydrogels spanning physiological and supraphysiological stiffnesses (1, 5, and 10 kPa), where substrate deformation provides a quantitative measure of cellular traction forces (Fig. 2A–C).

**Figure 2.**
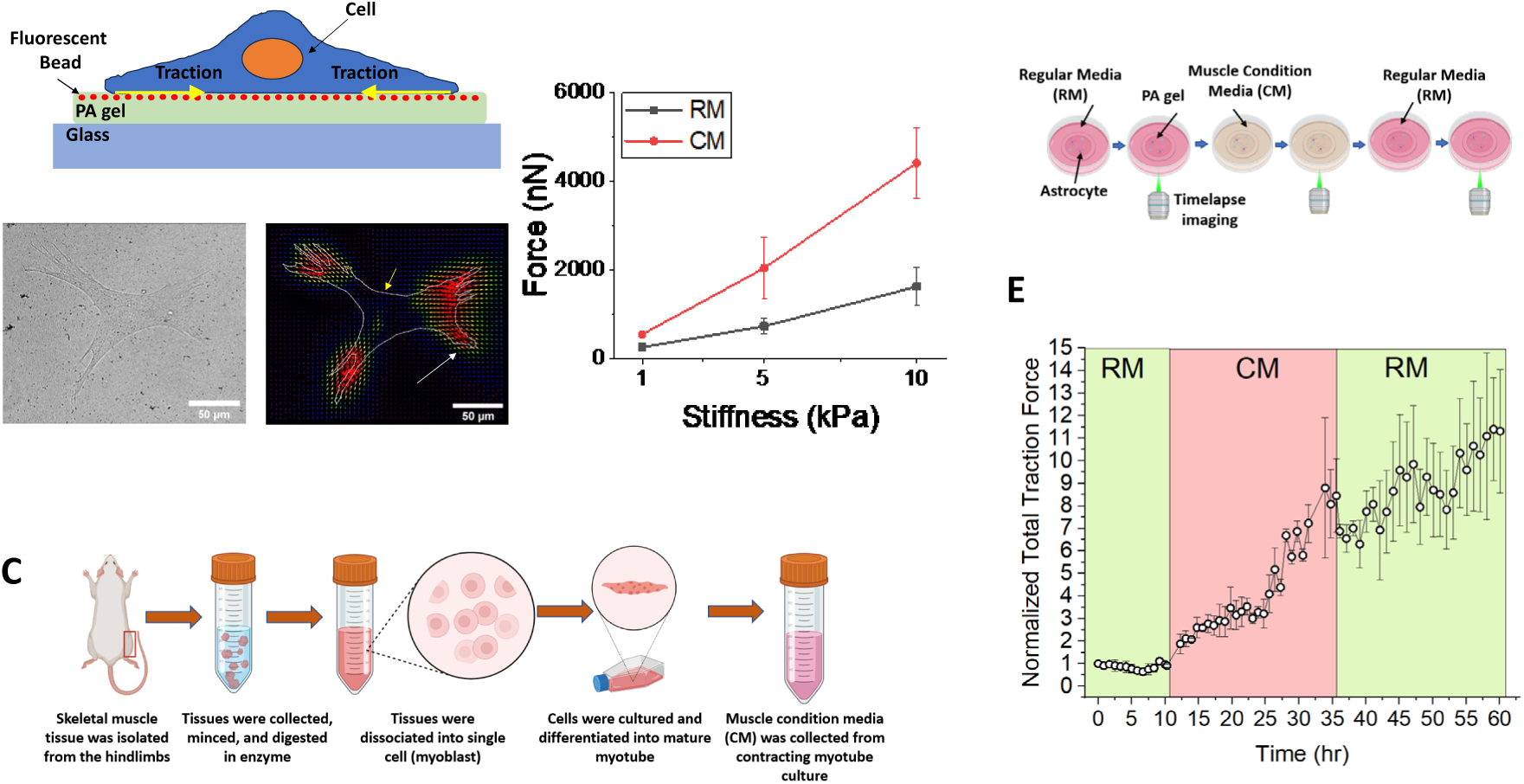
Muscle-conditioned media (CM) enhances astrocyte traction force in a stiffness-dependent manner. (A) Schematic and representative traction force microscopy (TFM) images illustrating astrocyte-induced substrate deformation on polyacrylamide (PA) gels embedded with fluorescent beads. (B) Quantification of astrocyte traction force on substrates of varying stiffness (1, 5, and 10 kPa) shows CM significantly increases force generation compared to regular media (RM) in a stiffness-dependent manner. Data represent mean ± SEM, n = 3 per group (C) Schematic of CM preparation: skeletal muscle was harvested from mouse hindlimbs, enzymatically dissociated into single myoblasts, differentiated into mature myotubes, and CM was collected from the contracting muscle culture. (D) Experimental setup for time-lapse TFM: astrocytes seeded on PA gels were first cultured in RM, then switched to CM, and imaged over time. (E) Time-course of total traction force exerted by astrocytes, normalized by their initial force, showing a marked increase after CM treatment. Astrocytes also maintain elevated force levels for at least 24 hours post-CM exposure. Data represent mean ± SEM, n = 3.

Under regular media (RM), astrocytes generated progressively greater traction forces on stiffer substrates, consistent with their intrinsic mechanical sensitivity (permutation test, *P* = 0.002; Fig. 2B). Exposure to CM for 24 h significantly increased traction force generation at all substrate stiffnesses, although the magnitude of the response depended on the mechanical environment. On 10 kPa substrates, CM increased traction forces approximately 4.5-fold (∼1000 to ∼4500 nN), whereas increases were approximately 2.5-fold on 5 kPa substrates (∼600 to ∼1500 nN) and 1.3-fold on 1 kPa substrates (∼150 to ∼200 nN) (Fig. 2B). Permutation analysis confirmed significant effects of media (*P* = 0.0005) and substrate stiffness (*P* = 0.0001), together with a significant media × stiffness interaction (*P* = 0.01), indicating that the contractile response to CM is strongly influenced by the mechanical environment.

We next examined the temporal dynamics of this response using live-cell TFM on astrocytes cultured on 5 kPa substrates. Baseline traction forces were recorded under RM for 10 h before switching to CM for 24 h, followed by a return to RM for an additional 24 h (Fig. 2E). CM induced a progressive increase in traction force within individual astrocytes, reaching approximately eightfold above baseline after 24 h.

Following CM withdrawal, force generation continued to increase, ultimately reaching approximately elevenfold above baseline after an additional 24 h in RM. Thus, transient exposure to muscle-derived factors initiated a sustained contractile response that persisted after removal of the stimulus.

Together, these findings demonstrate that hippocampal astrocytes generate substantial traction forces that are regulated by extracellular matrix stiffness and are markedly enhanced by exercise-associated muscle-derived signals. The persistence of elevated force generation following CM withdrawal suggests that these signals induce a sustained mechanically activated astrocyte state.

### Astrocyte contraction drives proliferation and growth

Having established that muscle-derived signals induce a sustained mechanically activated astrocyte state, we next asked whether astrocyte mechanical contractility is required for the proliferative and structural responses to these signals. Primary hippocampal astrocytes were cultured on polyacrylamide hydrogels of defined stiffness (1, 5, or 10 kPa) and treated with regular media (RM) or CM. Astrocyte number was quantified by DAPI and GFAP labeling on days 1 and 3 (Figs. 2A,C). No differences were observed on day 1. By day 3, CM induced a strong stiffness-dependent increase in astrocyte number, with ∼10-fold and ∼14-fold increases on 5 kPa and 10 kPa substrates, respectively, relative to RM, while a modest increase was detected on 1 kPa substrates (Fig. 3C). Significant main effects of media (CM > RM; p < 0.0002) and stiffness (p < 0.0002), as well as a media × stiffness interaction (p < 0.0002) from a permutation analysis, supported this conclusion. These results indicate that CM-driven astrocyte proliferation emerges only under conditions that support effective force generation, rather than as a simple biochemical response.

**Figure 3.**
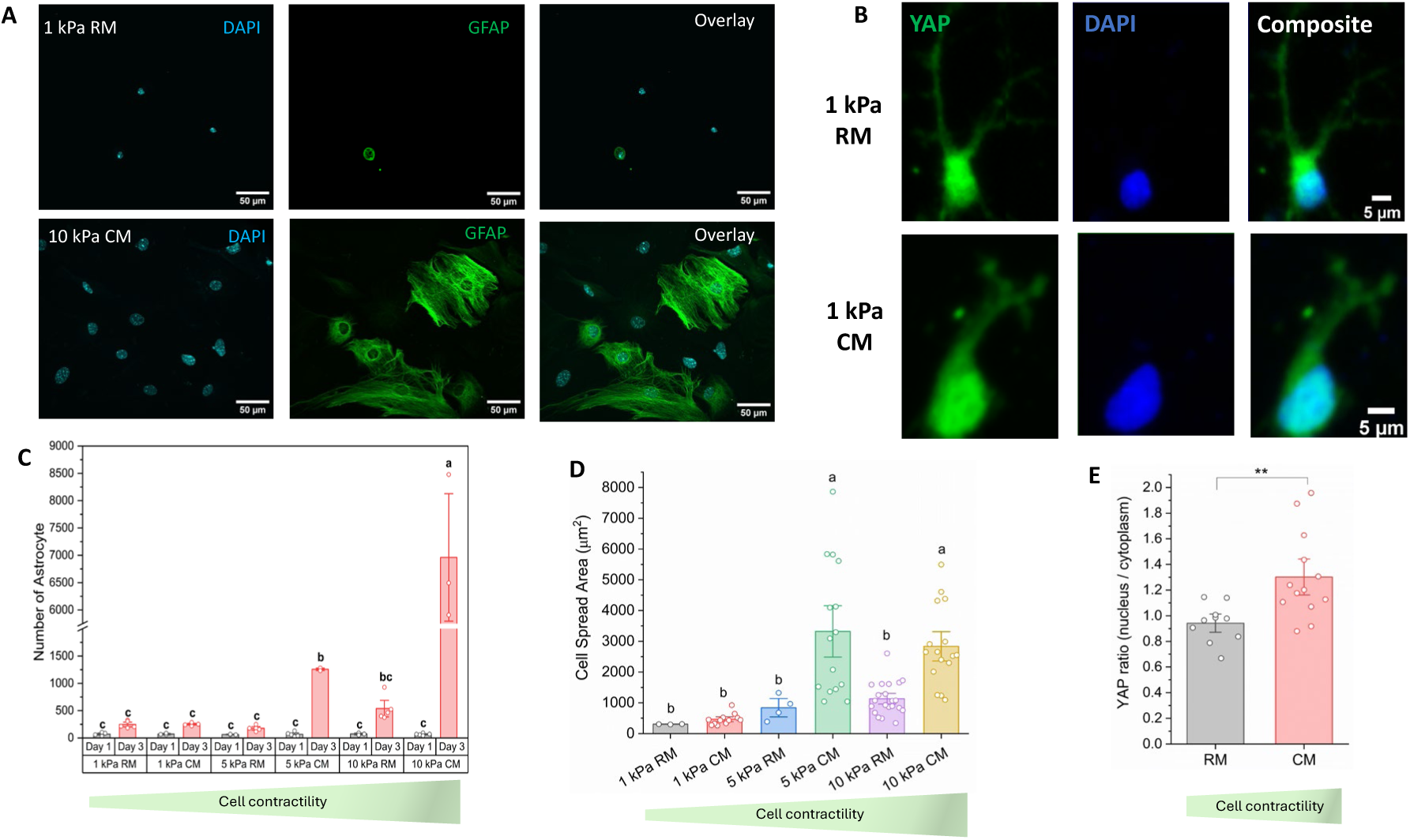
Mechanocontractility drives astrocyte phenotype and function: proliferation, cell spread area, and YAP nuclear translocation *in vitro*. (A) Representative immunofluorescence images of hippocampal astrocytes cultured on polyacrylamide (PA) gels with different stiffnesses (shown: 1 kPa RM and 10 kPa CM). Cells were stained with DAPI (nuclei, blue) and GFAP (astrocytes, green). CM treatment promoted greater astrocyte number and spreading area particularly on stiffer substrates. Scale bar: 50 μm. (B) Representative immunofluorescence images of YAP (green) and DAPI (blue) in astrocytes cultured on 1 kPa PA gel and treated with regular media (RM) or CM for 30 hours. CM treatment leads to enhanced nuclear localization of YAP. (C) Quantification of astrocyte number at Day 1 and Day 3. No significant proliferation was observed at Day 1 across groups. By Day 3, CM induced a dramatic increase in astrocyte proliferation in a stiffness-dependent manner, with the highest cell counts on 10 kPa CM. Data represent mean ± SEM, n = 3 per bar. Trapezoid below shows increasing substrate stiffness and cell force from left to right qualitatively (see Fig. 2 B). (D) Quantification of astrocyte spread area at Day 3 showed significantly larger spreading under CM treatment, especially on stiffer substrates. Data represent mean ± SEM, n = 3-21 per bar. Trapezoid below shows increasing substrate stiffness and cell force from left to right qualitatively (see Fig. 2 B). (E) Quantification of YAP nuclear-to-cytoplasmic ratio indicates a significant increase in CM-treated astrocytes compared to those in RM. Astrocytes were cultured on 1 kPa PA gel substrate. Data represent mean ± SEM, n = 10-13 per group. Trapezoid below shows cell force from left to right qualitatively.

To determine whether proliferation scales directly with force magnitude, we compared these outcomes with traction force microscopy measurements (Fig. 2 B). In RM, astrocyte traction forces increased ∼10-fold between 1 and 10 kPa, whereas proliferation increased only ∼2-fold (Fig. 3C) across the same range. Importantly, astrocytes cultured on 5 kPa CM substrates generated forces comparable to those on 10 kPa RM substrates yet exhibited substantially greater proliferation. This dissociation demonstrates that while astrocyte contractility is required to enable proliferation, CM provides additional signals that act in parallel with force rather than merely increasing it.

Finally, we asked whether CM-induced proliferation is accompanied by structural remodeling at the single-cell level. Quantification of astrocyte spread area revealed a pronounced, stiffness-dependent increase in cell size following CM treatment. On 5 and 10 kPa substrates, CM increased astrocyte area by ∼2.5–3.5-fold relative to RM, whereas only a modest increase (∼1.4-fold) was observed on soft 1 kPa gels (Fig. 3D). This was supported by significant main effects of media (CM > RM; F_1,65_=26.0, p < 0.0001) and stiffness (10 > 5 > 1 kPa; F_2,65_= 16.0, p < 0.0001) by 2-way ANOVA. The interaction between media × stiffness was marginally not significant (F_2,65_= 2.6, p=0.09). Together, these data demonstrate that astrocyte mechanical contractility is sufficient to promote astrocyte proliferation and growth and is required for the full pro-proliferative effects of muscle-derived biochemical signals.

### Muscle-derived signals promote astrocyte YAP activation

To determine whether the contractile response to muscle-derived signals is accompanied by mechanical signaling in the astrocytes, we quantified nuclear localization of YAP. Astrocytes were cultured on compliant polyacrylamide substrates (1 kPa) to allow YAP activation to be evaluated under a mechanically permissive baseline condition.

Exposure to conditioned media (CM) for 24 h significantly increased the astrocyte nuclear-to-cytoplasmic YAP ratio compared with regular media (RM) (unpaired *t*-test, *t*_21_ = 3.1, *P* = 0.005; Fig. 3A,B). Thus, muscle-derived biochemical signals not only increased astrocyte traction force generation but also activated a downstream mechanical signaling response consistent with engagement of YAP-dependent transcription.

### Mechanically activated astrocytes relay muscle-derived signals to neuronal networks

To determine whether muscle-derived signals influence neuronal mechanics directly or indirectly through astrocytes, we developed a sequential relay paradigm that separated astrocyte activation from neuronal force measurements (Fig. 4A). Primary astrocytes were first exposed to conditioned media (CM) under defined mechanical conditions, after which astrocyte-conditioned media (AST-CM) was transferred to glial-reduced hippocampal neuronal cultures maintained on an ultrasensitive three-dimensional force sensor. This approach isolated neuron-specific mechanical responses downstream of astrocyte activation while eliminating direct contributions from astrocyte-generated forces.

**Figure 4.**
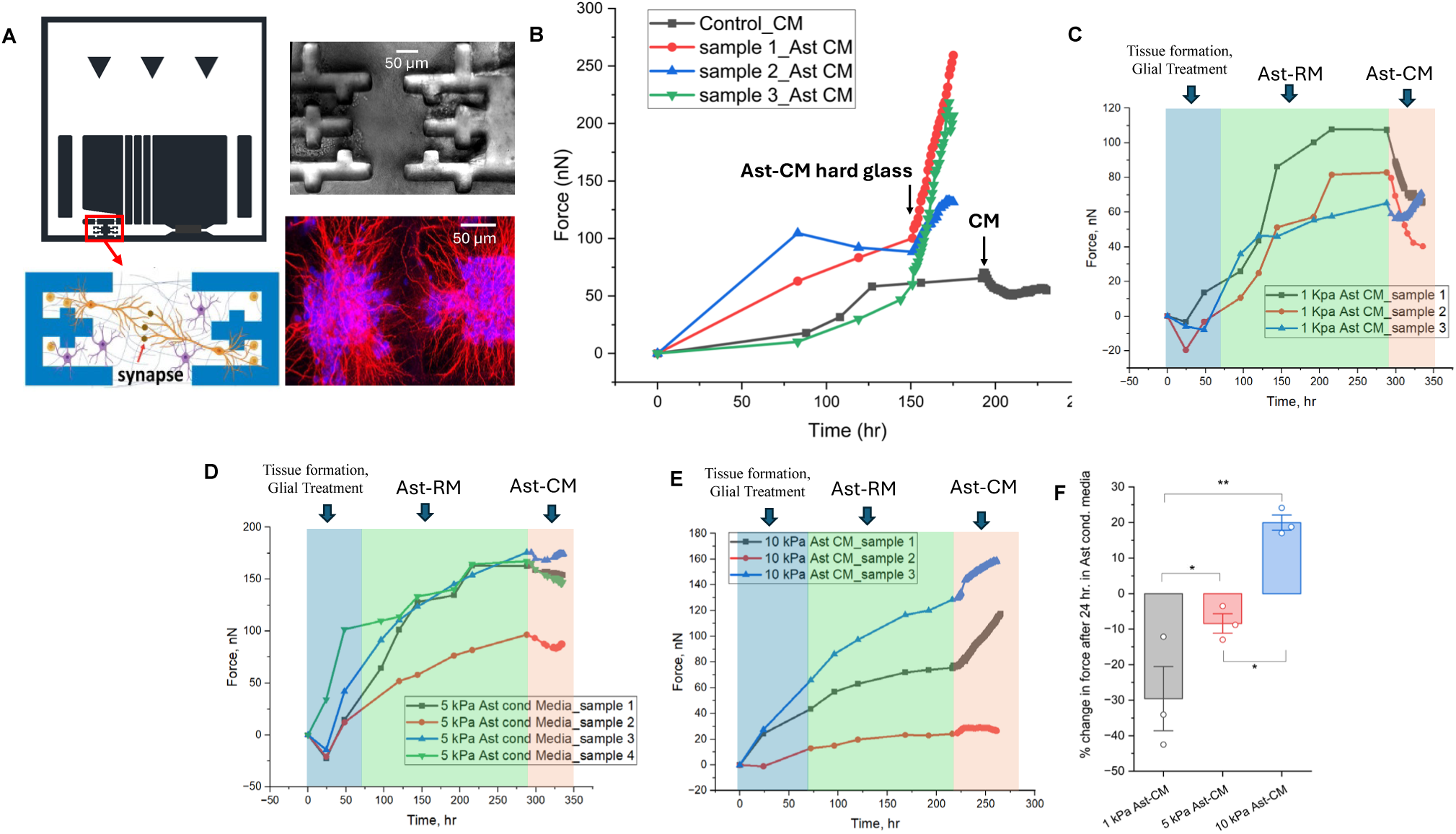
Astrocyte mechanotransduction relays muscle contraction to neuronal network contractility. (A) Schematic of the ultra-sensitive force sensor used to measure contractile forces generated by three-dimensional (3D) neuronal tissues. The illustration depicts neuronal growth on the sensor, where neuronal cell bodies are placed within opposing grips, extend neurites toward the central region, and form interconnected synaptic networks. A representative bright-field image of a 3D neuronal tissue formed on the force sensor, and stained image of neurons (MAP2: red, DAPI: blue) are shown on the right. Scale bar: 50 μm. (B) Representative force evolution of neuronal tissues exposed to control conditioned media (CM) or activated astrocyte-conditioned media (Ast-CM). At time zero the primary hippocampal neuronal cell cultures were exposed to a glial treatment to remove glial cells and hence isolate neurons (see methods for protocol). At that time, they also received a 1:1 mixture of neurobasal media with media from astrocytes cultured on hard glass substrate (Ast-RM). At approximately 150 hrs, media in three replicate samples were switched to Ast-CM with astrocytes cultured on hard glass. All three samples showed pronounced increase in neuronal force that progressively increased for at least 24 hrs. In one additional sample, at approximately 200 hrs, Ast-RM was switched with 1:1 mixture of neurobasal media with CM, and a slight decrease in force was observed. (C–E) Time-dependent tissue force generation for tissues treated with astrocyte-conditioned media derived from astrocytes cultured on **1 kPa** (C), **5 kPa** (D), and **10 kPa** (E) PA gels. Shaded regions denote the time duration of various treatments. Individual traces represent force of individual samples. (F) Percent change in tissue force 24 h after Ast-CM exposure, normalized to the force immediately before Ast-CM, demonstrating a stiffness-dependent response with reduced force following 1 kPa and 5 kPa activated astrocyte-conditioned media and enhanced force following 10 kPa activated astrocyte-conditioned media. Data are shown as mean ± SD with individual data points overlaid.

Glial-reduced neuronal cultures maintained in baseline astrocyte-conditioned media developed progressively increasing contractile forces during network maturation, consistent with previous observations (*23*) (Fig. 4B). Direct application of CM to these cultures at day 7 in vitro did not significantly alter neuronal contractility (Fig. 4B), indicating that muscle-derived factors alone were insufficient to increase neuronal network tension under these conditions.

We next examined whether astrocyte activation was required to relay the CM signal. Primary astrocytes were exposed to CM for 24 h, washed extensively to remove residual CM, and subsequently maintained in basal media to collect AST-CM. Because astrocytes remained mechanically activated following CM withdrawal (Fig. 2E), the collected media contained factors released from activated astrocytes without direct contamination by CM. Application of this AST-CM to glial-reduced neuronal cultures produced a robust increase in neuronal contractile force (Fig. 4B), indicating that mechanically activated astrocytes release soluble signals sufficient to enhance neuronal network tension.

To determine whether astrocyte mechanical state regulates this signaling response, astrocytes were cultured on substrates of defined stiffness (1, 5, or 10 kPa) before CM exposure and AST-CM collection. AST-CM derived from astrocytes cultured on 1 kPa substrates decreased neuronal contractile force, whereas AST-CM from 10 kPa substrates produced a marked increase in neuronal tension, with intermediate effects observed at 5 kPa (Figs. 4C–F). Across three independent experiments, the percent change in neuronal contractility 24 h after AST-CM treatment differed significantly among substrate stiffnesses (one-way ANOVA, *F*₂,₆ = 19.8, *P* = 0.002), with all pairwise comparisons reaching significance (*P* < 0.05).

Together, these findings support a model in which CM acts indirectly on neurons by initiating a mechanically gated astrocyte relay: CM induces astrocyte contractility, and the magnitude of astrocyte force generation, set by the mechanical environment, determines the composition and functional impact of astrocyte-secreted signals on neuronal networks. This mechanobiological pathway provides a means by which peripheral muscle activity can influence central neuronal mechanics and network behavior.

### Mechanically activated astrocytes promote immature neuron formation

Having established that muscle-derived signals induce a mechanically activated astrocyte state and alter neuronal network mechanics, we next asked whether this response influences the abundance of immature neurons. Because astrocytes are key regulators of adult hippocampal neurogenesis (*31*, *32*), we quantified doublecortin-positive (DCX⁺) immature neurons (*45*) following exposure to conditioned media collected from astrocytes maintained under defined mechanical conditions.

Primary astrocytes were cultured on polyacrylamide substrates spanning 1–10 kPa and exposed to either conditioned media (CM) or regular media (RM). After 24 h, CM was removed and astrocytes were maintained in basal media for an additional 24 h before collection of astrocyte-conditioned media (AST-CM). Conditioned media were subsequently applied to primary hippocampal cultures, and DCX⁺ immature neurons were quantified on day 9 (Fig. 5A,B).

**Figure 5.**
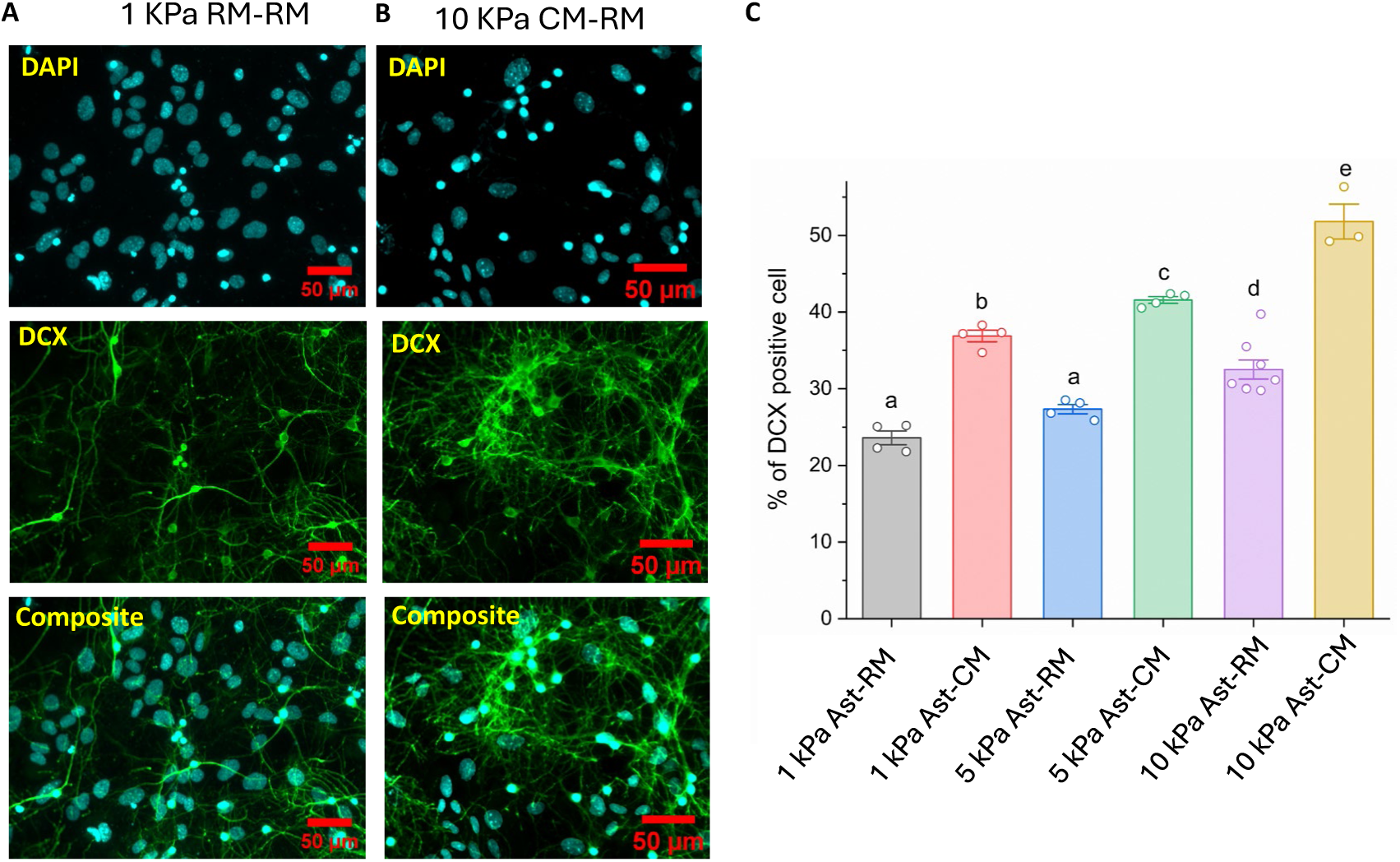
Muscle-to-astrocyte mechanotransduction generates a neurogenic secretome that enhances hippocampal neurogenesis. (A) Representative high-magnification immunofluorescence images of hippocampal neurons treated with astrocyte condition media, (Ast-CM), from 1 kPa substrates, stained for DAPI (cyan) and DCX (green). Scale bar: 50 μm. (B) Representative high-magnification immunofluorescence images of hippocampal neurons treated with astrocyte condition media, Ast-CM, from 10 kPa substrates, stained for DAPI (cyan) and DCX (green). Scale bar: 50 μm. (C) Quantification of DCX-positive cells shows that both substrate stiffness and Ast-CM exposure significantly increase the proportion of immature neurons, with the highest neurogenesis observed in the 10 kPa Ast-CM group. Bars with different letters denote statistically significant differences (p < 0.05). Data represent mean ± SEM, n = 3-8 culture dishes per group.

Media collected from astrocytes cultured on stiffer substrates consistently produced greater numbers of DCX⁺ immature neurons than media collected from astrocytes cultured on compliant substrates (Fig. 5C), indicating that the astrocyte mechanical state influences the release of pro-neurogenic signals. This effect was observed under both RM and CM conditions. However, CM further enhanced DCX⁺ neuron abundance beyond that associated with substrate stiffness alone. In particular, CM-treated astrocytes cultured on 1 or 5 kPa substrates promoted greater DCX⁺ neuron formation than RM-treated astrocytes exhibiting comparable levels of contractility, indicating that muscle-derived signals contribute additional pro-neurogenic activity beyond their effects on astrocyte mechanics. The largest increase in DCX⁺ neurons occurred in the 10 kPa CM condition, where elevated contractility and CM stimulation coincided. Two-way ANOVA confirmed significant main effects of media (*P* < 0.0001) and substrate stiffness (*P* < 0.0001), together with a significant interaction between the two variables (*P* = 0.004).

Together, these findings indicate that the mechanical state of astrocytes regulates their capacity to promote immature neuron formation. Muscle-derived signals further augment this response through mechanisms that are not fully explained by increased contractility alone, supporting a model in which biomechanical and biochemical signals converge to regulate astrocyte-mediated neurogenic signaling.

Collectively, our findings define a mechanically activated astrocyte state, enhanced by muscle-derived signals, and characterized by increased actomyosin activity, mechanical signaling, structural remodeling, and altered paracrine function. This coordinated response provides a mechanistic framework linking exercise-derived muscle signals to hippocampal plasticity.

## DISCUSSION

Physical exercise is among the most effective stimuli promoting hippocampal plasticity, yet the cellular mechanisms linking peripheral muscle activity to coordinated changes within the brain have remained incompletely understood. Here, we identify a mechanically activated astrocyte state that is engaged during exercise and is associated with changes in astrocyte signaling, neuronal mechanics, and immature neuron formation. In vivo, voluntary running was accompanied by rapid activation of astrocyte mechanical signaling together with progressive recruitment of actomyosin machinery. Using complementary in vitro models, we show that muscle-derived biochemical signals induce sustained astrocyte force generation, which contributes to astrocyte proliferation, cellular expansion, and YAP nuclear translocation, while altering the release of astrocyte-derived factors that regulate neuronal network tension and the abundance of DCX⁺ immature neurons. Together, these findings support a model in which regulated astrocyte mechanics contribute to communication between peripheral muscle activity and hippocampal plasticity.

Voluntary exercise produced two spatially and temporally distinct responses in hippocampal hilar astrocytes. Astrocytic YAP localization was tightly correlated with locomotor activity during the preceding 1–2 minutes, whereas pMLC was most strongly associated with cumulative running over the preceding 30–40 minutes. The rapid YAP response was observed selectively in the hilus and was absent in the molecular layer, indicating that exercise engages astrocyte mechanotransductive signaling in a spatially restricted manner. Although YAP can also be regulated by non-mechanical pathways (*18*), the minute-scale, behavior-locked dynamics observed here closely resemble mechanically driven YAP responses and are difficult to reconcile with slower biochemical signaling cascades alone (*18*, *41*, *42*, *46*). This interpretation is reinforced by the complementary pMLC response. Whereas YAP provides a rapid readout of mechanotransductive signaling, phosphorylation of myosin light chain reflects activation of the non-muscle myosin II contractile apparatus (*43*, *44*) that may take 30-40 mins to de-phosphorylate. Together, these distinct temporal profiles suggest that astrocytes rapidly engage mechanical signaling during locomotion while the contractile machinery is progressively recruited over tens of minutes of sustained running.

Rather than functioning solely as passive mechanical sensors of their local environment our findings suggest that astrocytes actively integrate biochemical signals released from exercising muscle with the local mechanical environment to generate a distinct mechanical and secretory state. Unlike previous studies emphasizing how astrocytes respond to externally imposed mechanical cues (*19*), our results indicate that physiological exercise induces astrocytes to become active mechanical effectors whose contractile state influences neighboring neural cells. Consistent with this model, mechanically activated astrocytes released factors that increased neuronal network mechanics and promoted the abundance of DCX⁺ immature neurons, whereas muscle-conditioned media alone had little effect on neuronal mechanics in glial-reduced cultures. Moreover, the magnitude and direction of these responses depended on the mechanical state of the astrocytes from which the conditioned media was obtained. Together, these findings suggest that astrocytes function as mechanically regulated intermediaries that translate peripheral muscle-derived signals into local paracrine signals capable of remodeling the neurogenic niche (*47*). One potential mechanism is that contractility alters vesicle trafficking, membrane tension, exocytosis, or extracellular vesicle release, thereby modifying the composition or amount of secreted signaling molecules, although identifying these mediators will require future study (*48–53*).

These findings also provide a new perspective on the relationship between mechanics and adult hippocampal neurogenesis. Astrocytes have long been recognized as essential regulators of the neurogenic niche through the secretion of trophic and gliotransmitter molecules (*31*, *32*). Importantly, the stiffness of the substrate where astrocytes were cultured on and muscle-conditioned media each increased DCX+ cells independently, indicating that astrocyte mechanics alone do not fully account for the observed neurogenic effects. Instead, our data support a model in which mechanical activation creates a permissive state that is subsequently shaped by exercise-associated biochemical signals to regulate astrocyte secretory activity and downstream neurogenic responses. This framework expands the role of astrocytes from passive recipients of mechanical information to active regulators of tissue mechanics and neural plasticity.

Several limitations should be considered. First, the in vivo experiments measured mechanical signaling and activation of the contractile machinery rather than force generation directly. Second, although the in vitro studies establish that muscle-derived factors are sufficient to induce a mechanically activated astrocyte state, they do not identify the molecular components responsible for this response. Finally, while our data support a model in which astrocyte mechanics influence neuronal function and immature neuron formation, additional loss-of-function studies targeting astrocyte contractility will be required to establish the necessity of this pathway in vivo.

More broadly, our findings suggest that mechanics represents an important dimension of intercellular communication in the nervous system. Mechanical forces influence membrane organization, cytoskeletal dynamics, vesicle trafficking, and transcriptional regulation throughout biology (*23*, *54–58*), yet these processes are rarely considered within the framework of exercise-induced brain plasticity. By integrating mechanical and biochemical signaling, astrocytes may provide a means through which peripheral physiological activity is translated into coordinated cellular responses within the hippocampus. More generally, the stiffness dependence of astrocyte contractility observed here suggests one potential cellular mechanism by which changes in brain mechanical properties could influence astrocyte function, perhaps contributing to reported associations between regional brain stiffness and cognitive performance (*59–64*). Future development of genetically encoded tension reporters and in vivo force imaging approaches (*65*) will be essential for determining how astrocyte mechanics are regulated during behavior and how they contribute to neural circuit function.

Collectively, our findings support a model in which exercise engages a mechanically activated astrocyte state that integrates peripheral muscle-derived signals with local tissue mechanics to regulate astrocyte signaling, neuronal mechanics, and immature neuron formation. Incorporating regulated cellular force generation into current models of brain plasticity broadens the conceptual framework through which astrocyte function and exercise-induced hippocampal remodeling are understood.

## Acknowledgments

Research reported in this publication was supported by the National Science Foundation (NSF) under award numbers 2342257 (to MTAS and JR), 2123781 (Mind In-Vitro) (to MTAS), and Chan Zuckerberg Biohub (Chicago) Investigator grant (to MTAS). The content is solely the responsibility of the authors and does not necessarily represent the official views of NSF or Chan Zuckerberg Biohub.

## Author contributions

MSHJ and MTAS conceived and designed *in vitro* experiments

IEM, MGC, and JSR conceived and designed *in vivo* experiments

MSHJ, IEM and MGC performed the experiments.

IEM, MGC, JPR, JS, YD performed *in vivo* confocal imaging

JSR conducted statistical analysis

MGC and IEM conducted mouse brain slicing

MSHJ, MGC and IEM conducted immunostaining

IEM, JS, DL, JPR, and NS conducted IMARIS image analysis of *in vivo* data

MSHJ prepared force sensor and PA gel

QVG prepared force sensors and PA gel, helped with image analysis

MA prepared force sensors and PA gel, helped with image analysis

MAZ and YH prepared force sensors and PA gel

MSHJ and KYL collected muscle condition media MSHJ and BE prepared force sensor mold

MSHJ, MTAS, JSR and IEM analyzed and interpreted data and analysis

MSHJ wrote the first draft of the manuscript

MTAS and JSR reviewed and edited the manuscript.

Funding acquisition: MTAS and JSR

## Data, code, and materials availability

All data and code needed to evaluate and reproduce the results in the paper are present in the paper. This study did not generate any new materials.

